# A dual-target herbicidal inhibitor of lysine biosynthesis

**DOI:** 10.1101/2022.03.04.482975

**Authors:** Emily R. R. Mackie, Andrew S. Barrow, Rebecca M. Christoff, Belinda M. Abbott, Anthony R. Gendall, Tatiana P. Soares da Costa

**Affiliations:** Department of Biochemistry and Genetics, La Trobe Institute for Molecular Science, La Trobe University, Bundoora, VIC 3086, Australia; School of Agriculture, Food and Wine and Waite Research Institute, University of Adelaide, Waite Campus, Glen Osmond, SA 5064, Australia; Department of Chemistry and Physics, La Trobe Institute for Molecular Science, La Trobe University, Bundoora, VIC 3086, Australia; Australian Research Council Industrial Transformation Research Hub for Medicinal Agriculture, AgriBio, La Trobe University, Bundoora, VIC 3086, Australia; Department of Animal, Plant and Soil Sciences, La Trobe University, Bundoora, VIC 3086, Australia

## Abstract

Herbicides with novel modes of action are urgently needed to safeguard global agricultural industries against the damaging effects of herbicide-resistant weeds. We recently developed the first herbicidal inhibitors of lysine biosynthesis, which provided proof-of-concept for a promising novel herbicide target (Soares da Costa et al., 2021). In this study, we expanded upon our understanding of the mode of action of herbicidal lysine biosynthesis inhibitors. We previously postulated that these inhibitors may act as proherbicides (Soares da Costa et al., 2021). Here we show this is not the case. We report an additional mode of action of these inhibitors, through their inhibition of a second lysine biosynthesis enzyme, and investigate the molecular determinants of inhibition. Furthermore, we extend our herbicidal activity analyses to include a weed species of global significance.

## INTRODUCTION

Effective herbicides are critical for sustainable agriculture. However, our current options are dwindling as the prevalence of herbicide-resistant weeds continues to rise (Hall et al., 2020). Weeds have now evolved resistance to 21 out of the 31 herbicide modes of action, yet there has been a lack of herbicides with new modes of action brought to market over the last 30 years (Duke, 2011; Heap, 2022).

Despite the success of targeting amino acid biosynthesis enzymes for the development of herbicides (e.g., glyphosate and chlorsulfuron), the inhibition of plant lysine biosynthesis has never been explored commercially. Our group was the first to develop inhibitors of lysine biosynthesis with herbicidal activity (Soares da Costa et al., 2021). We showed that the most potent of these inhibitors, (*Z*)-2-(5-(4-methoxybenzylidene)-2,4-dioxothiazolidin-3-yl)acetic acid (MBDTA-2) (Figure 1A) targets lysine production by inhibiting dihydrodipicolinate synthase (DHDPS), the enzyme that catalyses the first and rate-limiting step in the pathway (Soares da Costa et al., 2018). Interestingly, we found that the mode of DHDPS inhibition by MBDTA-2 was through binding at a novel allosteric site distinct from the allosteric lysine binding site (Figure 1B), which enables regulation of the enzyme (Hall et al., 2021).

**Figure 1.**
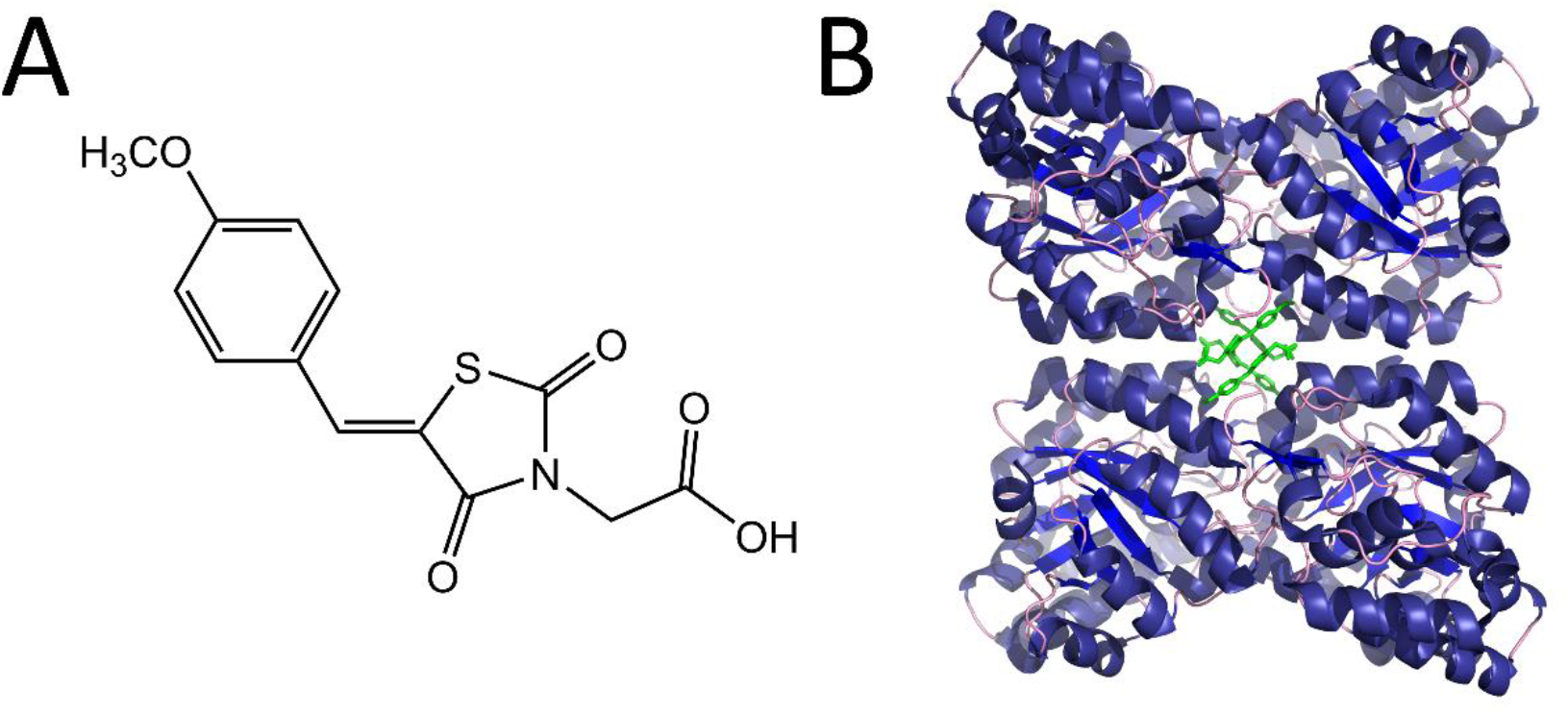
Structure and mode of binding of MBDTA-2. (A) Chemical structure of MBDTA-2. (B) Cartoon view of the AtDHDPS1 quaternary structure with MBDTA-2 (green sticks) bound within a novel allosteric pocket (PDB ID: 7MDS) (Soares da Costa et al., 2021).

In the previous study, we revealed that the *in vitro* potency of MBDTA-2 against recombinant *Arabidopsis thaliana* (At) DHDPS enzymes was similar to the activity against agar-grown *A. thaliana* (Soares da Costa et al., 2021). Usually, herbicides inhibit their enzyme targets with greater potency than they inhibit *in vivo* plant growth. This is because the amount of herbicide reaching the target site is less than the amount applied. We proposed that this unusual similarity between *in vitro* and *in vivo* activity may be due to MBDTA-2 acting as a proherbicide, which is modified *in vivo* to a form that is more active at the target site. Proherbicides have been reported in the literature such as diclofop-methyl, which is demethylated via ester hydrolysis *in vivo* to produce the more active compound diclofop (Shimabukuro et al., 2002). Whilst MBDTA-2 could not undergo the same process as it does not contain an ester, we postulated that a similar proherbicidal effect may be observed through the *in vivo* demethylation of the aryl methyl ether.

The present study sought to extend our understanding of the mode of action of our previously developed DHDPS inhibitors (Christoff et al., 2021; Soares da Costa et al., 2021). Specifically, we used biochemical enzyme kinetic assays to demonstrate that MBDTA-2 does not act as a proherbicide, and that the apparent similarity between *in vitro* and *in vivo* potency may instead be explained by this series of compounds having a novel, dual-target mode of action through inhibition of the second lysine biosynthesis enzyme in the pathway, dihydrodipicolinate reductase (DHDPR). Static docking and biochemical assays revealed that in contrast to the allosteric binding of these inhibitors to DHDPS, active site binding is responsible for their inhibition of DHDPR. Additionally, we successfully extended our previous herbicidal activity studies on *A. thaliana* to include one of the most agriculturally problematic weeds in the world, rigid ryegrass (*Lolium rigidum*) (Bajwa et al., 2021; Busi and Beckie, 2021).

## RESULTS

### Inhibitory activity of a demethylated MBDTA analogue

Given that proherbicides are metabolised *in vivo* to produce compounds with greater potency at the target site, it can be assumed that the metabolised form will have greater activity than the proherbicidal form against the target *in vitro*. As such, to assess whether MBDTA-2 is a proherbicide that is demethylated *in vivo*, we measured the inhibitory activity of the demethylated analogue, (*Z*)-2-(5-(4-hydroxybenzylidene)-2,4-dioxothiazolidin-3-yl)acetic acid (HBDTA), against both recombinant *A. thaliana* DHDPS enzymes (Figure 2) (Christoff et al., 2021). The dose-response curves yielded IC_50_ values for AtDHDPS1 and AtDHDPS2 of 100 ± 0.95 μM and 105 ± 1.04 μM, respectively (Figure 2). These values are slightly greater than those we reported for MBDTA-2 (IC_50_ (AtDHDPS1) = 63.3 ± 1.80 μM, IC_50_ (AtDHDPS2) = 64.0 ± 1.00 μM) (Soares da Costa et al., 2021). This data suggests, conversely to our hypothesis, that the activity of MBDTA-2 at the target site is not influenced by the retention or loss of the methyl group *in vivo*.

**Figure 2.**
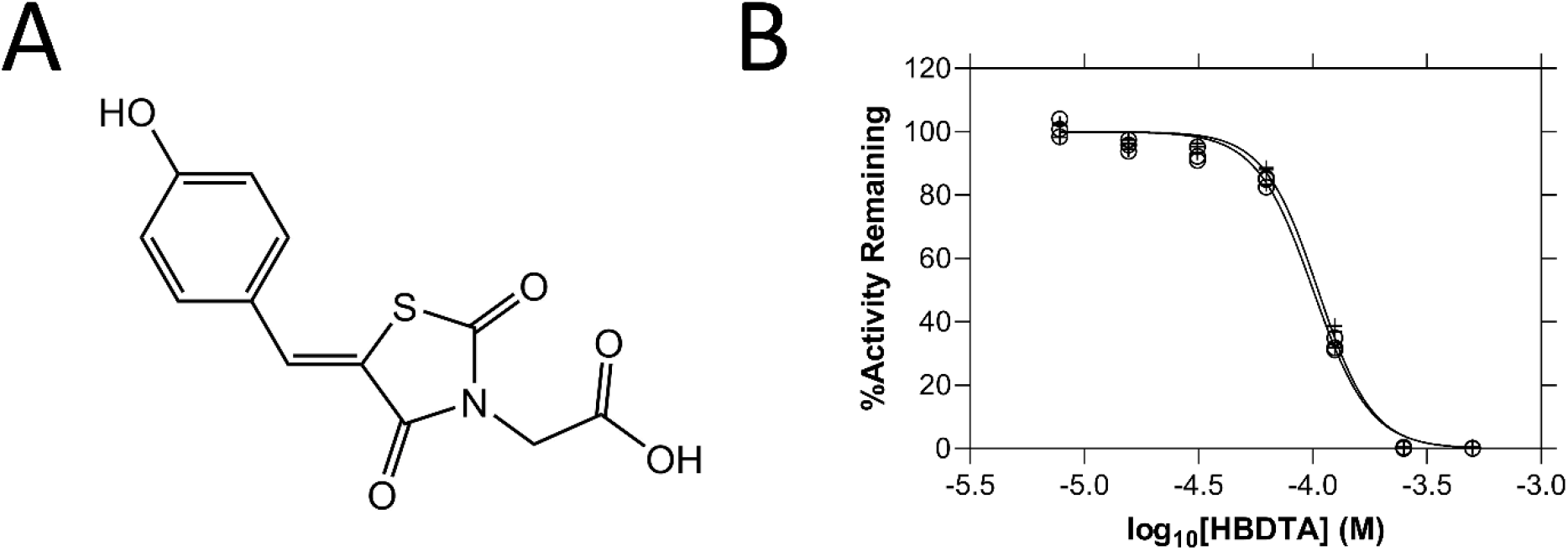
Structure and *in vitro* potency of HBDTA. (A) Chemical structure of HBDTA. (B) Dose-response curves of HBDTA against recombinant AtDHDPS1 (○) and AtDHDPS2 (+) enzymes. Initial enzyme rate was normalised against the vehicle control to determine % activity remaining. Data were fitted to a nonlinear regression model (solid line), resulting in R^2^ values of 0.99.

### Dual-target activity of MBDTA-2

Given that the similarity in the *in vitro* and *in vivo* activity of MBDTA-2 could not be explained by enhanced target site activity of the demethylated compound, we sought to explore other mechanisms that may explain this observation. We investigated whether additional modes of action beyond the inhibition of the DHDPS enzyme may account for the increased *in vivo* potency relative to the *in vitro* activity against the target enzyme. It is well established that proteins catalysing sequential reactions within a metabolic pathway often have conserved binding site features (Hsu et al., 2013; Jensen, 1976; Jenwitheesuk et al., 2008; Zhang et al., 2009). As such, we hypothesised that DHDPS inhibitors may also have activity against the enzyme following DHDPS in the plant lysine biosynthesis pathway, DHDPR. The activity of both recombinant *Arabidopsis thaliana* DHDPR enzymes was measured whilst titrating MBDTA-2 to determine the IC_50_ values of 6.92 ± 0.92 μM against AtDHDPR1 and 8.58 ± 1.19 μM against AtDHDPR2 (Figure 3). These results demonstrate that MBDTA-2 is a multi-targeted inhibitor of two consecutive enzymes in the lysine biosynthesis pathway, AtDHDPS and AtDHDPR. This compound represents the first example of a dual-target inhibitor of the lysine biosynthesis pathway.

**Figure 3.**
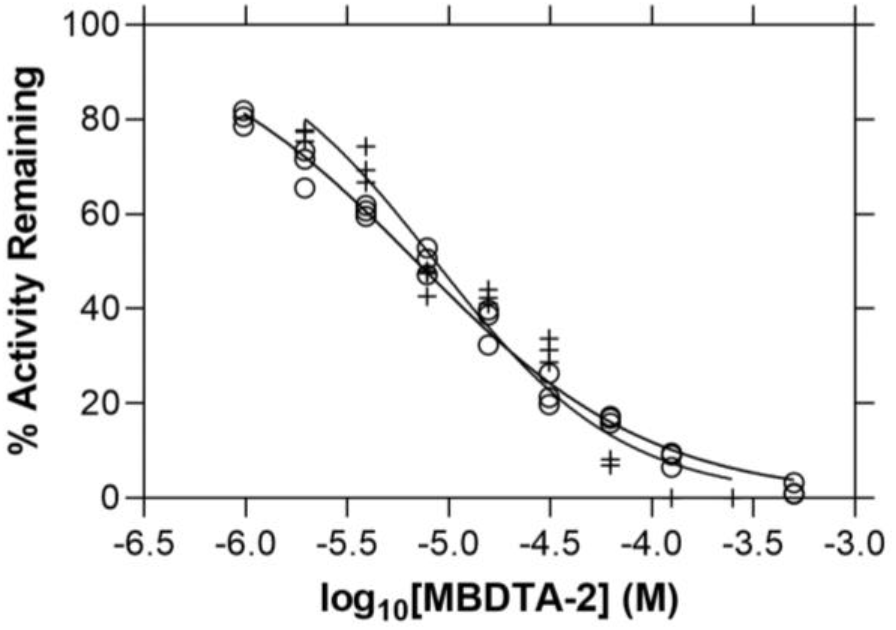
*In vitro* potency of MBDTA-2 against AtDHDPR. Dose-response curves of MBDTA-2 against recombinant AtDHDPR1 (○) and AtDHDPR2 (+) enzymes. Initial enzyme rate was normalised against the vehicle control to determine % activity remaining. Data were fitted to a nonlinear regression model (solid line), resulting in R^2^ values of 0.99 and 0.95 for AtDHDPR1 and AtDHDPR2, respectively.

### Mode of AtDHDPR inhibition by MBDTA-2

To investigate the molecular determinants of inhibition of AtDHDPR, we sought to co-crystallise the enzyme with MBDTA-2. Given that our attempts were unsuccessful, we employed a static docking approach using the published AtDHDPR2 crystal structure (Watkin et al., 2018). The resulting data suggested that MBDTA-2 binds in the active site (Figure 4A). The hydrophobic pocket occupied by MBDTA-2 overlaps with the probable NADPH cofactor binding site, based on the crystal structure of cofactor-bound *Escherichia coli* DHDPR (Reddy et al., 1996; Scapin et al., 1997). The predicted MBDTA-2 orientation suggests its stabilisation by polar interactions between the heterocyclic ring and Thr122 and Gly120. Additionally, the MBDTA-2 acid is within hydrogen bonding proximity to Asp185. To validate the mechanism of inhibition of MBDTA-2 against AtDHDPR, further enzyme kinetic experiments were performed. The previous dose-response experiments were conducted according to standard practice in that substrate and cofactor were kept at limiting concentrations to ensure that inhibition may be measured regardless of the kinetic mechanism of inhibition. Subsequently, the activity of AtDHDPR was measured whilst titrating MBDTA-2, this time in the presence of excess amounts of substrate and nucleotide cofactor, i.e., at concentrations 10-fold above the respective *K_M_* values (Figure 4B). The IC_50_ values determined were 72.7 ± 1.07 μM and 69.5 ± 1.06 for AtDHDPR1 and AtDHDPR, respectively, which are 10-fold and 8-fold greater than those determined for AtDHDPR1 and AtDHDPR2 when substrate and cofactor were limiting. This apparent reduction in potency indicates that MBDTA-2 is a competitive inhibitor, and therefore likely binds at the AtDHDPR active site as suggested by the docking results. Interestingly, this contrasts with the allosteric mode of inhibition of MBDTA-2 against AtDHDPS.

**Figure 4.**
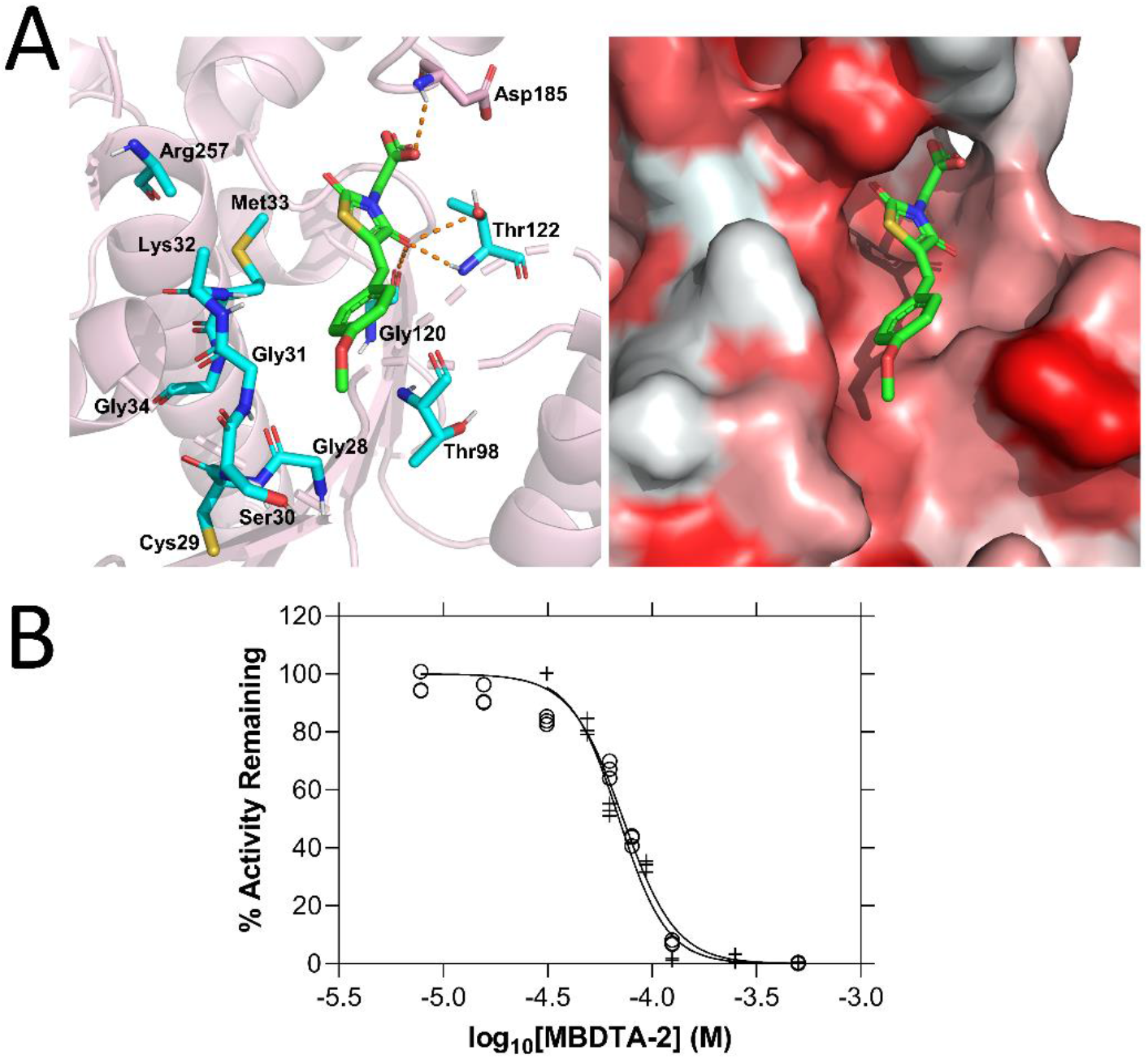
Mode of AtDHDPR2 inhibition by MBDTA-2. (A) The predicted MBDTA-2 (green) binding site resulting from static docking with AtDHDPR2 (PDB ID: 5UA0) overlaps with the probable NADPH cofactor binding site (cyan, left panel). Hydrophobicity of the predicted binding pocket (right panel) is represented by white-red shading indicating hydrophilic-hydrophobic residues. (B) Dose-response curves of MBDTA-2 against AtDHDPR1 (○) and AtDHDPR2 (+) enzymes in the presence of saturating concentrations of substrate and cofactor. Data were fitted to a nonlinear regression model (solid line), resulting in R^2^ values of 0.97 and 0.98 for AtDHDPR1 and AtDHDPR2, respectively.

### Herbicidal activity of MBDTA-2 against weeds

Previously, we showed that the MBDTA-2 compound has herbicidal activity against the model plant *A. thaliana* and is therefore the first example of a herbicidal lysine biosynthesis inhibitor. To further assess the potential of inhibiting plant lysine biosynthesis enzymes for the development of herbicides, the efficacy of MBDTA-2 against the economically significant invasive weed species rigid ryegrass *L. rigidum* was investigated. Treatment of *L. rigidum* with 1200 mg·L^−1^ (equivalent to 48 kg·ha^−1^) of MBDTA-2 resulted in inhibition of plant germination and growth, corresponding to a significant reduction in shoot fresh and dry weight and a significant reduction in root dry weight (Figure 5). Specifically, we observed ~4-fold and ~5-fold reductions in shoot fresh and dry weight, respectively, and a ~2-fold reduction in root dry weight (Figure 5). These results further exemplify the potential of lysine biosynthesis inhibitors for development as herbicide candidates, as we have demonstrated that MBDTA-2 possesses herbicidal activity against one of the most problematic weed species to global agriculture.

**Figure 5.**
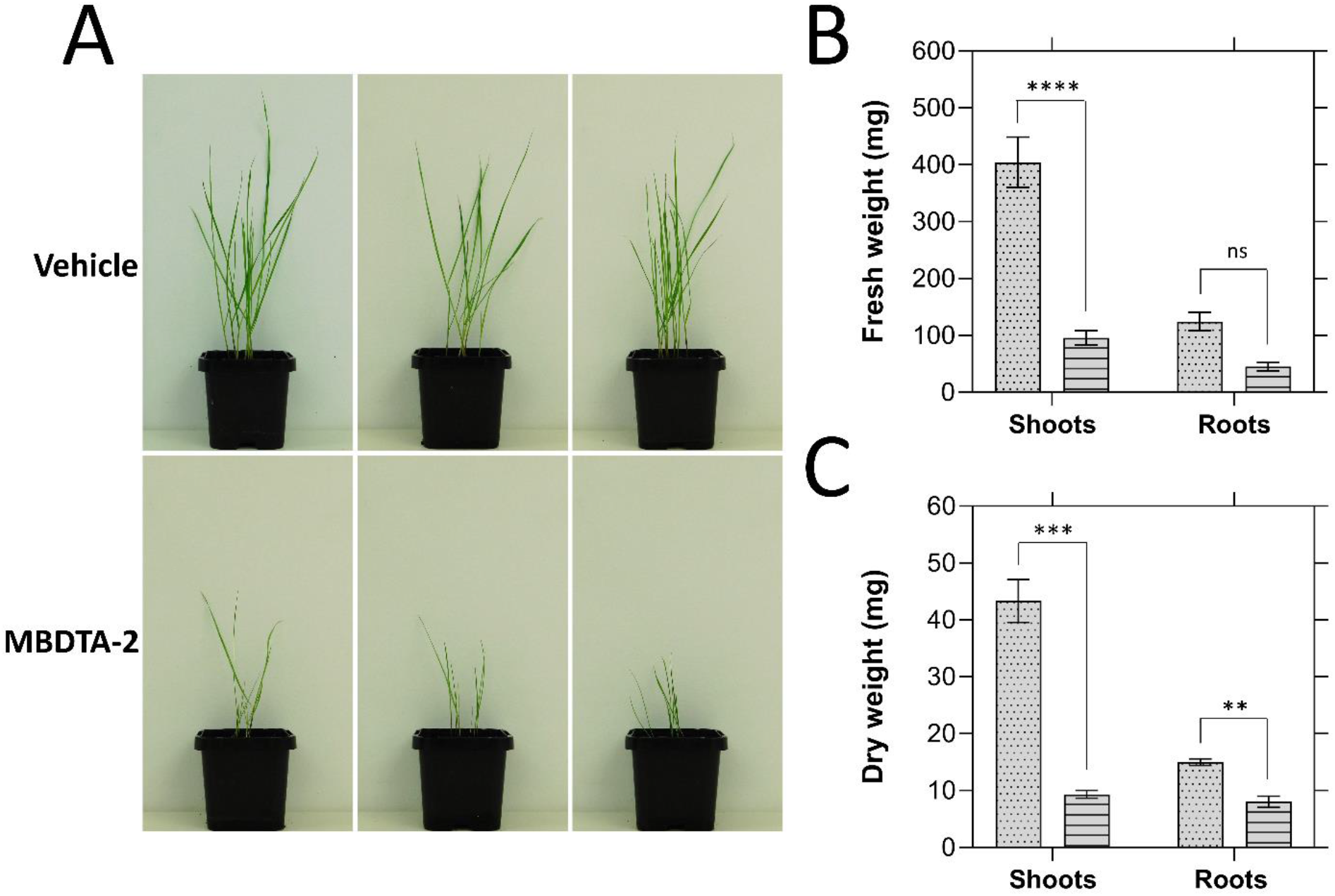
Inhibition of *Lolium rigidum* germination and growth by MBDTA-2. (A) 14-day growth of *L. rigidum* treated with three pre-emergence treatments of vehicle control (2% (v/v) DMSO, 0.01% Agral) or 1200 mg·L^−1^ of MBDTA-2. (B) Fresh weight of *L. rigidum* shoots and roots following treatment of plants with vehicle control (dots) or MBDTA-2 (lines). Shoots, *P* = 0.00002, roots, *P* = 0.05233, unpaired Student’s two-tailed Etest. (C) Dry weight of *L. rigidum* shoots and roots following treatment of plants with vehicle control (dots) or MBDTA-2 (lines). Shoots, *P* = 0.00088, roots, *P* = 0.00374, unpaired Student’s two-tailed *t*- test. Data were normalised against the vehicle control. Data represents mean ± S.E.M. (*N* = 3). \*\**P* < 0.01, \*\*\**P* < 0.001, \*\*\*\**P* < 0.0001.

## DISCUSSION

Herbicides with new modes of action are urgently needed to combat the rise in herbicide-resistant weed species, which pose a global threat to agricultural industries. In our previous study, we described the development of the first herbicidal lysine biosynthesis inhibitors, providing proof-of-concept for a novel herbicide mode of action. Whilst we previously explored the molecular mode of action of these inhibitors at their target site, namely the DHDPS enzyme, an explanation for their unusual similarity in *in vitro* and *in vivo* potency remained to be delineated. Specifically, we hypothesised that they may be acting as proherbicides, through demethylation of their aryl methyl ethers. While this mechanism of proherbicide conversion has not been reported to date, the demethylation of aryl methyl ethers on prodrugs, such as codeine, is well established (Dayer et al., 1988; Kirchheiner et al., 2006). Nevertheless, our finding that the demethylated analogue of the MBDTA-2 aryl methyl ether did not impact activity against the target enzyme DHDPS demonstrated that these compounds are not proherbicides.

Given that we could not attribute the similarity between the *in vitro* and *in vivo* potency of our compounds to their modification to a more active form *in vivo*, we postulated that we may have previously failed to capture the totality of their target site effects. Our discovery that MBDTA-2 is an inhibitor of not only DHDPS, but also of the subsequent enzyme in the lysine biosynthesis pathway DHDPR, supported this hypothesis. Moreover, the ~8-fold greater potency of MBDTA-2 against DHDPR than DHDPS reveals that the *in vitro* potency is actually ~6-fold greater than the *in vivo* potency. The phenomenon of inhibitors having dual-target activity against consecutive enzymes in metabolic pathways has sometimes been attributed to conserved active site features (Hsu et al., 2013; Toulouse et al., 2020). However, there are also many examples of inhibitors of multiple targets from distinct pathways, which have been identified regardless of binding site similarities (Allen et al., 2015; Hashmi et al., 2021; Wang et al., 2019). Nevertheless, to our knowledge, this is the first time a dual-target inhibitor has been shown to have allosteric and orthosteric inhibitory activity at two different enzymes. The advantages of dual-target inhibitors over single-target inhibitors have been well-reported for the development of novel drugs (Allen et al., 2015; Oldfield and Feng, 2014). Such advantages include a reduced susceptibility to the generation of resistance, which is also a highly desirable property in herbicide development (Gressel, 2020; Hall et al., 2020). Indeed, there has been a focus on the use of herbicide mixtures inhibiting multiple molecular targets in attempts to reduce the generation of resistance to existing herbicides (Gressel, 2020). Despite the recognition of the potential of inhibiting multiple targets for the reduction of target site resistance generation, little work has been done on the development of new herbicides that do so (Fu et al., 2019; Giberti et al., 2017). Dual-target herbicidal compounds such as MBDTA-2 are therefore promising candidates for progressing the herbicide development field beyond the ‘one target-one herbicide’ approach.

The development of new herbicides that are effective against *L. rigidum*, particularly those with a reduced propensity to generate resistance, is of the highest priority given the economic impact of this species. An overreliance on a small number of herbicide modes of action has culminated in the widespread evolution of multiple resistance mechanisms in *L. rigidum* (Bajwa et al., 2021; Owen et al., 2014). In Australia alone, herbicide-resistant *L. rigidum* invades 8 million hectares of cropping land, resulting in revenue losses of AUD$93 million annually (Llewellyn et al., 2016). Our finding that MBDTA-2 can significantly reduce *L. rigidum* germination and growth further illustrates the potential utility of lysine biosynthesis inhibitors in combatting the global herbicide resistance crisis.

## MATERIALS AND METHODS

### Chemical synthesis

Compounds were synthesised as previously described (Christoff et al., 2021; Perugini et al., 2018).

### Protein expression and purification

Recombinant AtDHDPS1, AtDHDPS2, AtDHDPR1 and AtDHDPR2 proteins were produced as previously described (Mackie et al., 2022; Soares da Costa et al., 2021).

### Enzyme inhibition assays

DHDPS enzyme activity was measured using methods previously described (Soares da Costa et al., 2021). DHDPR enzyme activity was measured using methods previously described (Mackie et al., 2022). Briefly, reaction mixtures were incubated at 30 °C for 12 min before a second 60 s incubation following the addition of excess *E. coli* DHDPS (51 μg·mL^−1^) for generation of the DHDP substrate. The relevant DHDPR isoform (2.6 μg·mL^−1^) was added to initiate the reaction, and substrate turnover measured spectrophotometrically at 340 nm via the associated oxidation of the cofactor NADPH.

### Docking

The AtDHDPR2 crystal structure was retrieved from the Protein Data Bank and hydrogens added using AutoDock Tools. Three dimensional MBDTA-2 was docked with AtDHDPR2 using an unlimited search space in the PyRX interface using AutoDock Vina with default parameters. The resulting ligand poses were visualised in PyMol.

### Herbicidal activity analyses

The herbicidal efficacy of MBDTA-2 against *L. rigidum* was assessed using methods similar to those reported previously (Mackie et al., 2022). Pre-wet seed-raising soil (pH 5.5) (Biogro, Dandenong South, VIC, Australia) supplemented with 0.22% (w/w) Nutricote N12 Micro 140 day-controlled release fertiliser (Yates, Sydney, NSW, Australia) was used. 10 seeds were sown at a depth of 0.5 cm into pots of pre-wet soil, following stratification at 4 °C for 21 days in the dark. Compounds dissolved in DMSO were diluted to working concentrations in H_2_O containing 0.01% (v/v) Agral (Syngenta, North Ryde, NSW, Australia) to a final DMSO concentration of 2% (v/v). Treatments were given by pipetting 2.0 mL of MBDTA-2, vehicle control or positive control (chlorosulfuron PESTANAL®(Sigma-Aldrich, North Ryde, NSW, Australia)) directly onto seeds upon sowing and on each of the subsequent two days. Plants were grown in a chamber at 22 °C under a 16 h light (100 μmol m^−2^ s^−1^)/8 h dark schedule for 14 days before photos were taken. Roots and shoots were separated prior to drying at 70 °C for 72 h. Experiments were performed in biological triplicates.

## ACKNOWLEDGEMENTS

T.P.S.C. acknowledges the Australian Research Council for funding support through a DECRA Fellowship (DE190100806). Work in A.R.G.’s laboratory is supported by the Australian Research Council Research Hub for Medicinal Agriculture (IH180100006). E.R.R.M. acknowledges the Grains Research and Development Corporation (9176977) for support through a PhD scholarship and operational funding and La Trobe University for support through a Research Training Program Scholarship. R.M.C. is a recipient of an Australian Government Research Training Program Scholarship and a LIMS Write-Up Award.

## AUTHOR CONTRIBUTIONS

E.R.R.M.: investigation, formal analysis, data curation, visualisation, methodology, validation, writing – original draft preparation, writing – review and editing

A.S.B.: investigation, formal analysis, data curation, methodology, validation, writing – review and editing

R.M.C.: investigation, formal analysis, data curation, methodology, validation, writing – review and editing

B.M.A.: resources, supervision, writing – review and editing

A.R.G.: resources, supervision, writing – review and editing

T.P.S.C.: conceptualisation, resources, supervision, funding acquisition, writing – review and editing

